# Deep Learning for Protein Structure Prediction: Advancements in Structural Bioinformatics

**DOI:** 10.1101/2023.04.26.538026

**Authors:** Daniel Szelogowski

## Abstract

**Motivation:** Accurate prediction of protein structures is crucial for understanding protein function, stability, and interactions, with far-reaching implications in drug discovery and protein engineering. As the fields of structural bioinformatics and artificial intelligence continue to converge, a standardized model for protein structure prediction is still yet to be seen as even large models like AlphaFold continue to change architectures. To this end, we provide a comprehensive literature review highlighting the latest advancements and challenges in deep learning-based structure prediction, as well as a benchmark system for structure prediction and visualization of amino acid protein sequences.

**Results:** We present ProteiNN, a Transformer-based model for end-to-end single-sequence protein structure prediction, motivated by the need for accurate and efficient methods to decipher protein structures and their roles in biological processes and a system to perform prediction on user-input protein sequences. The model leverages the transformer architecture’s powerful representation learning capabilities to predict protein secondary and tertiary structures directly from integer-encoded amino acid sequences. Our results demonstrate that ProteiNN is effective in predicting secondary structures, though further improvements are necessary to enhance the model’s performance in predicting higher-level structures. This work thus showcases the potential of transformer-based architectures in structure prediction and lays the foundation for future research in structural bioinformatics and related fields.

## 1 Introduction

End-to-end single-sequence protein structure prediction is a task in Bioinformatics to predict the three-dimensional (3D) structure of a protein from its amino acid sequence. In an end-to-end single-sequence model, the input is a protein sequence (as a string of amino acids, typically then integer encoded), and the output is a predicted 3D structure in a standard format such as **PDB (Protein Data Bank)**. The model is trained to learn the relationship between the amino acid sequence and the nascent 3D structure of the protein, often either by angles or coordinates. The main advantage of single-sequence prediction is that it does not require any additional information about the protein other than its amino acid sequence, unlike traditional methods that often rely on additional information such as homology information or predicted secondary structure. Single-sequence prediction is still challenging, however; its accuracy is not always sufficient for practical applications, so it is often used in combination with other methods or as a preliminary step in more complex structure prediction pipelines.

Protein structure prediction is currently being applied in numerous fields:

- **Drug discovery** — accurate predictions of protein structures can aid researchers in the design of new drugs that target specific proteins. By understanding the structure of a protein, researchers can identify potential binding sites for drugs, which can be helpful in the design of new drugs.
- **Biotechnology** — predicted structures can be applied to design new enzymes and other biotechnology products that are more efficient and effective.
- **Disease diagnosis** — understanding the structure of proteins involved in diseases can help researchers better understand disease mechanisms and develop new therapies. For example, predictions of the structures of proteins involved in cancer can be used to design new drugs that target these proteins.
- **Agriculture** — predictions can be used to improve crop yields and develop new crops that are more resistant to pests and diseases.
- **Environmental monitoring** — predictions can be used to monitor ecological pollutants and understand their effects on living organisms.

The task of structure prediction presents numerous challenges for researchers — although recent advances in deep learning have provided a valuable baseline for accelerating research on the topic. A significant degree of variability in protein structures makes it difficult to predict the 3D structure from the amino acid sequence. Protein structures are also dynamic, meaning they can change over time, making it challenging to capture the full range of structures in a single prediction. Neural Networks and other Deep Learning models have been able to capture the complex relationships between the amino acid sequence and the 3D structure of proteins, leading to more accurate predictions, as well as being able to handle the large amounts of data that are required for protein structure prediction — making them well-suited for this task.

We present ProteiNN, a Transformer-based model for end-to-end single-sequence protein structure prediction, designed to advance our understanding of protein folding and function. By leveraging transformer architectures’ powerful representation learning capabilities, ProteiNN predicts protein secondary and tertiary structures directly from integer-encoded amino acid sequences. We discuss the implementation, evaluation, and limitations of the ProteiNN model, explore its potential applications in structural bioinformatics, and provide a comprehensive review of the literature. This work demonstrates the utility of transformer-based models in predicting protein structures and paves the way for future advancements in computational biology, drug discovery, and protein engineering.

While significant advances such as AlphaFold and ESMFold have greatly improved the accuracy of structure prediction, they rely heavily on large datasets, Multiple Sequence Alignments, and substantial computational resources. In contrast, we propose a lightweight, interpretable transformer-based model (ProteiNN) intended not to compete with state-of-the-art methods, but to serve as a reproducible, educational benchmark for single-sequence structure prediction. This dual-purpose work offers both a literature review contextualizing the field and a hands-on system to illustrate modern deep learning approaches to protein folding, especially for secondary structure prediction.

## 2 Related Work

Deep learning has revolutionized the field of protein structure prediction, driving significant advancements in understanding complex biological systems. Various neural network architectures — such as **Convolutional Neural Networks (CNNs), Recurrent Neural Networks (RNNs)**, and **Recurrent Geometric Networks (RGNs)** — have been employed to tackle the challenges associated with predicting protein structures. These innovative approaches have enabled researchers to model complex protein structures more accurately, unlocking new insights into protein function, interaction, and evolution. This literature review explores the application of these deep learning architectures in the realm of structure prediction, highlighting their respective strengths, limitations, and potential for future advancements in the field.

### 2.1 Convolutional Neural Networks

Yang et al. investigate the efficiency and effectiveness of using CNNs in place of Transformer-based models for pre-trained protein sequence language models [1]. Current protein language models are limited in scalability due to the quadratic scaling of Transformers, which restricts the maximum sequence length that can be analyzed. The authors thus introduce **CARP (Convolutional Autoencoding Representations of Proteins)**, a CNN-based architecture that scales linearly with sequence length. The study demonstrates that CARP models are competitive with the state-of-the-art Transformer model ESM-1b [2] across various downstream applications, including structure prediction, zero-shot mutation effect prediction, and out-of-domain generalization. The study challenges the association between masked language modeling and Transformers, highlighting that the pre-training task, not the Transformer architecture, is essential for making pre-training effective. Furthermore, CARP shows strong performance on sequences longer than those allowed by current Transformer models, suggesting that computational efficiency can be improved without sacrificing performance using a CNN architecture.

### 2.2 Recurrent Neural Networks

Torrisi et al. review the recent advancements in protein structure prediction that have been bolstered by the introduction of Deep Learning techniques, including the adoption of RNNs, **Long Short-Term Memory (LSTM)** networks, and **Bidirectional RNNs (BRNNs)** [3]. These models have excelled at handling sequential data and learning long-range dependencies, making them particularly well-suited for protein sequence analysis. In recent years, various protein structure predictors have been developed, such as SPOT-Contact, which combines CNNs and 2D BRNNs with LSTM units to improve the accuracy of contact map prediction. By leveraging these recurrent architectures, researchers have been able to exploit evolutionary information, yielding more sophisticated pipelines for protein structure prediction tasks.

While the literature has seen an upsurge in methods utilizing recurrent neural models, other Deep Learning approaches such as CNNs, **Feed-Forward Neural Networks (FFNNs)**, and **Residual Networks (ResNets)** have also made significant contributions to the field. Methods such as DeepCDpred, DeepContact, DeepCov, DNCON2, MetaPSICOV, PconsC4, RaptorX-Contact, TripletRes, and AlphaFold have each employed unique architectures and input features to improve protein structure prediction. These advancements have resulted in considerable improvements in contact and distance map predictions, directly impacting the quality of 3D protein structure predictions. As computational resources, novel techniques, and experimental data continue to grow, further progress in protein structure prediction is expected, with recurrent architectures playing a significant role in this rapidly advancing field.

### 2.3 Recurrent Geometric Networks

AlQuraishi introduces the novel end-to-end differentiable RGN architecture for protein structure learning [4]. This model aims to overcome the central challenge of predicting protein structures from sequences by coupling local and global structures using geometric units. The model achieved state-of-the-art accuracy in two challenging tasks — predicting novel folds without co-evolutionary data and known folds without structural templates. The RGN architecture allows a model to implicitly encode multi-scale protein representations and predict structures by integrating information from residues upstream and downstream. Likewise, RGNs are substantially faster than existing methods, potentially enabling new applications such as integrating structure prediction within docking and virtual screening.

RGNs learn an implicit representation of protein fold space using secondary structure as the dominant factor in shaping their representation despite not being explicitly encoded with the concept. The architecture can also complement existing methods, such as incorporating structural templates or co-evolutionary information as priors or inputs for learning to improve secondary structure prediction. As such, the author predicts that hybrid systems using deep learning and co-evolution as priors, along with physics-based approaches for refinement, will soon solve the long-standing problem of accurate and efficient structure prediction. However, the model has limitations, such as its reliance on **Position-Specific Scoring Matrices (PSSMs)**, which could potentially be addressed with more data-efficient model architectures. Our model thus draws inspiration from RGNs but focuses on a simplified architecture to prioritize interpretability and reproducibility over predictive performance.

### 2.5 Transformer Models & Attention Networks

Chandra et al. discuss the application of Transformer models from the field of **Natural Language Processing (NLP)** for predicting protein properties [5]. These models — referred to as protein language models — are capable of learning multi-purpose representations of proteins from large open repositories of protein sequences. The architecture has shown promising results in predicting protein characteristics such as post-translational modifications. It also provides advantages over traditional deep learning models, effectively capturing longrange dependencies in protein sequences (similar to natural language).

The authors review various protein prediction tasks for which Transformer models have been applied, including protein structure, protein residue contact, protein-protein interactions, drug-target interactions, and homology studies. These models have demonstrated impressive results without relying on **Multiple Sequence Alignments (MSAs)** or structural information. Additionally, the authors highlight the interpretability of Transformer models, which allows for visualization and analysis of attention weights and subsequently provides deeper biological insights.

While they have shown significant improvements over RNNs and other models, they possess limitations such as fixed-length input sequences and quadratic growth in memory requirements. Nevertheless, advancements are being made to address these issues, such as incorporating hidden states from previous fragments and implementing sparse attention mechanisms to reduce computational complexity. Although Transformers have outperformed other models in many tasks, they may not always be the best choice for all protein prediction tasks, and a combination of methods may be necessary. As the applications of Transformers in computational biology and bioinformatics are still in their infancy, further improvements and special-purpose models can be expected. The proof-of-principle example in this review displays the potential of using Transformers as general feature extractors to improve results compared to traditional protein features, like MSAs, but the authors note that more research is needed to determine their overall superiority. Ultimately, the future of computational biology and bioinformatics will likely involve advancements in Transformer models and their integration with other methods for protein analysis.

### 2.5 Language Models

Lin et al. demonstrate the potential of large language models for evolutionary-scale prediction of atomic-level protein structures [6]. By training models with up to 15 billion parameters, the authors discovered that the models’ understanding of protein sequences correlated with structure prediction accuracy. The study introduces **ESMFold**, a fully end-to-end single-sequence structure predictor, which achieved a speedup of up to 60x on the inference forward pass compared to state-of-the-art methods like AlphaFold and RosettaFold. They note that simplifying the neural architecture and eliminating the need for multiple sequence alignments contributed to the improvement in speed.

Similarly, the authors also present the ESM Metagenomic Atlas, which offers the first large-scale structural characterization of metagenomic proteins comprised of more than 617 million structures. This atlas provides an unprecedented view into the vast diversity of some of the least understood proteins on Earth. The combined results of ESMFold and the ESM Metagenomic Atlas indicate that as language models scale, their ability to predict protein structures improves, especially for proteins with low evolutionary depth. This advancement in speed and accuracy could accelerate the discovery of new protein structures and functions, leading to potential breakthroughs in medicine and biotechnology.

### 2.6 AlphaFold

Jumper et al. present a groundbreaking computational method capable of predicting protein structures with atomic accuracy even when no similar structure is known [7]. **AlphaFold**, the neural network-based model at the core of this work, has been completely redesigned to incorporate physical and biological knowledge about protein structure and multi-sequence alignments. In the 14th **Critical Assessment of [Protein] Structure Prediction (CASP14)**, AlphaFold demonstrated remarkable accuracy, outperforming other methods and achieving results competitive with experimental structures in a majority of cases.

AlphaFold’s success can be attributed to its novel neural architectures and training procedures, which are based on evolutionary, physical, and geometric constraints of protein structures. Key innovations include the use of a new architecture to jointly embed MSAs and pairwise features, a new output representation and associated loss for accurate end-to-end structure prediction, a new equivariant attention architecture, the use of intermediate losses for iterative refinement of predictions, masked MSA loss for joint training with the structure, and learning from unlabelled protein sequences through self-distillation and self-estimates of accuracy. These advances have enabled AlphaFold to significantly improve the accuracy of protein structure prediction and provide valuable insights into protein function and biology.

The end-to-end structure prediction in AlphaFold relies on a structure module that operates on a 3D backbone structure using the pair representation and the original sequence row from the MSA representation in the trunk. The structure module updates the global frame (residue gas) representation iteratively in two stages. First, it uses **Invariant Point Attention (IPA)** to update a set of neural activations without changing the 3D positions. Next, the updated activations perform an equivariant update operation on the residue gas. The **Frame-Aligned Point Error (FAPE)** (the final loss calculation) compares the predicted atom positions to the true positions under multiple alignments. Predictions of the structure’s side-chain angles and per-residue accuracy are computed with small per-residue networks at the end of the process. AlphaFold’s architecture is trained with both labeled and unlabeled data, enhancing accuracy using an approach similar to noisy student self-distillation. A separate structure module is trained for each of the 48 Evoformer blocks in the network, providing a trajectory of 192 intermediate structures that represent the network’s belief of the most likely structure at each block. The authors note that the accuracy of AlphaFold decreases when the median alignment depth is less than 30 sequences; however, the model is still effective for proteins with few intra-chain or homotypic contacts compared to the number of heterotypic contacts. The architectural concepts in AlphaFold are expected to apply to predicting full hetero-complexes in future systems, overcoming the difficulty with protein chains with many hetero-contacts.

## 3 Approach

Protein structure prediction aims to determine the 3D conformation of a protein given its amino acid sequence. Formally, let 𝒜 = {*A*_1_, *A*_2_, …, *A*_20_} be the set of 20 standard amino acids. A sequence of length *L* is represented as **s** = (*s*_1_, *s*_2_, …, *s*_L_) where *s*_*i*_ ∈ 𝒜. The goal is to learn a mapping *f* : 𝒜^*L*^ → ℝ^3*L*^ that predicts the 3D coordinates **X** = (**x**_1_, **x**_2_, …, **x**_L_) of the protein’s atoms.

The ProteinNet dataset is a comprehensive and standardized resource designed to facilitate the training and evaluation of data-driven models for protein sequence-structure relationships, including structure prediction and design [8]. The dataset integrates protein sequence, structure, and evolutionary information in machine learning-friendly file formats to provide accessibility for researchers. One of the critical components of ProteinNet is the high-quality **Multiple Sequence Alignments (MSAs)** of all structurally characterized proteins, which were generated using substantial high-performance computing resources. Additionally, the dataset includes standardized splits of data (e.g., train, train-eval, test, and validation) that emulate the difficulty of past CASP experiments by resetting protein sequence and structure space to the historical states preceding six prior CASPs.

To evaluate the quality of the predicted structures, we perform assessments using standard metrics including the **Root-Mean-Square Deviation (RMSD)** between the predicted and experimental structures (see *Section 5.1*). This metric provides a comprehensive view of the model’s ability to predict the overall fold, topological similarity, and atomic-level accuracy of protein structures. We also calculate classification metrics such as precision, recall, and *F*_1_ score (see *Section 5.2*).

## 4 Implementation

We present a Transformer-based model for end-to-end single-sequence protein structure prediction, implemented in Python 3.9 with the PyTorch library and trained on Windows 11 using an RTX 3060 Ti with 8GB of VRAM. The model accepts integer-encoded amino acid sequences and missing residue masks as input and predicts the dihedral angles, which are then used to generate PDB files representing the 3D structures. A user-friendly front-end application allows users to input protein sequences as strings, which are then encoded for the model [9].

### 4.1 ProteiNN

Our system features an attention-based model — **ProteiNN** — for predicting protein structure from amino acid sequences (see Figure 6). The model was trained and evaluated using the SideChain-Net dataset, which provides the basis for complete model training. Our approach predicts the 12 angles provided by the dataset, differing from prior models that only utilized 3 angles (see Figure 1). The model’s output is in the shape of *L*× 12× 2, with values between [− 1, 1], allowing us to handle the circular nature of angles more effectively [10].

**Figure 1:**
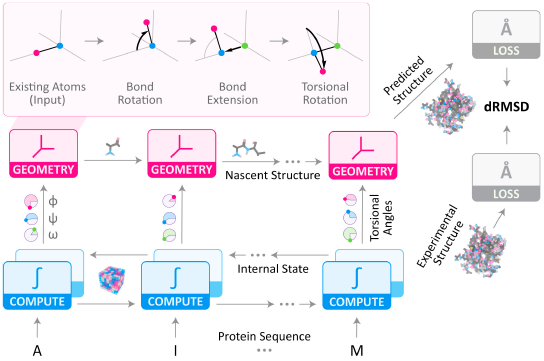
RGN architecture, modified for our system [4]

### 4.2 Model Architecture

Let **s** ∈ ℕ^*L*^ be the input sequence of length *L*, where each amino acid is encoded as an integer. The sequence is embedded into continuous representations:

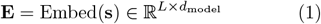

Our model employs a **Multi-Head Self-Attention** mechanism to capture the dependencies among amino acids and their spatial arrangement. The attention mechanism computes:

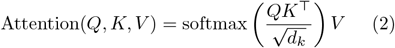

where *Q, K*, and *V* are the query, key, and value matrices obtained from the embedded input **E**:

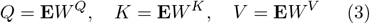

Here, *W*^*Q*^, *W*^*K*^, and *W*^*V*^ are learnable weight matrices, and *d*_*k*_ is the dimensionality of the key vectors [11].

For **Multi-Head Attention**, the mechanism is extended as:

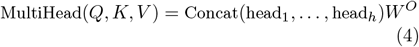

where each attention head is:

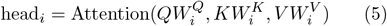

and 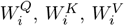,and *W*^*O*^ are learnable projection matrices.

### 4.3 Gating Mechanism

We incorporate a gating mechanism that modulates the information flow between the input and output:

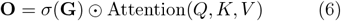

where *σ* is the sigmoid function, **G** is the gating vector computed from the input, and ⊙ denotes element-wise multiplication. This allows the model to selectively emphasize specific relationships and discard irrelevant information.

### 4.4 Masking for Missing Residues

To handle missing residue information, we introduce a mask **M** ∈ **{**0, 1**}**^*L*^, where *M*_*i*_ = 0 if residue *i* is missing and 1 otherwise. During the attention calculation, the masked positions are filled with a large negative value to effectively exclude them from the softmax normalization step, ensuring that the model ignores the missing residues:

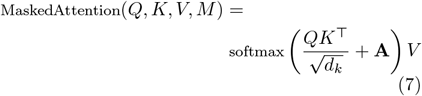

where **A** is the attention bias matrix defined as:

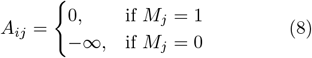

### 4.5 Angle Prediction and Loss Function

Because a simple Mean Squared Error (MSE) loss function does not account for the periodicity of angles (e.g., angles *π* and − *π* are equivalent), we predict the sine and cosine values for each angle. For each residue *i*, the model outputs 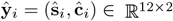,where ŝ_*i*_ and **ĉ**_*i*_ are the predicted sine and cosine values of the 12 dihedral angles.

The loss function is defined as the mean squared error between the predicted and true sine and cosine values:

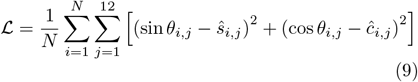

where *N* is the number of residues, *θ*_*i,j*_ is the true angle, and ŝ_*i,j*_, ĉ_*i,j*_ are the predicted sine and cosine values.

To recover the angles after they are predicted, we use the arctan 2 function:

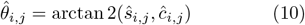

This ensures that mathematically equivalent angles (such as *π* and − *π*) are treated identically by the model, as periodicity ensures robustness against angular equivalence. This approach is a necessity in protein structure prediction, where angular periodicity is intrinsic to the data.

### 4.6 Geometric Reconstruction

The predicted angles are used to reconstruct the 3D coordinates of the protein using forward kinematics. Each dihedral angle defines the rotation around a bond in the protein backbone. The position of each atom can be computed using rotation matrices derived from the predicted angles.

Our system also provides methods for visualizing and comparing the predicted protein structures to the ground truth (see Figure 2). The **Batched-StructureBuilder** class from SideChainNet is used to generate these structures, taking a tensor of integers representing the amino acid sequences and a tensor of floating-point numbers representing the predicted angles for each residue.

**Figure 2:**
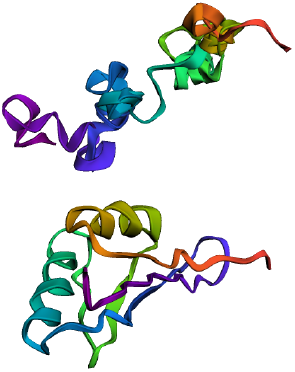
Example model prediction (top) and ground truth (bottom), visualized with Py3DMOL

## 5 Evaluation

We employ a rigorous evaluation framework to assess the performance of our Transformer-based protein structure prediction model, quantifying the accuracy of the model’s predictions using the RMSD, which estimates the overall structural differences between the predicted and experimental protein structures.

### 5.1 Root-Mean-Square Deviation (RMSD)

The RMSD between the predicted structure 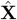 the true structure **X** is computed as:

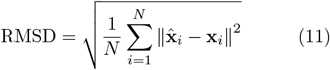

where *N* is the number of atoms, 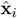 and **x**_*i*_ are the predicted and true coordinates of atom *i*, respectively. This metric provides a quantitative measure of the structural deviation at the atomic level.

Our model achieved a final training RMSE loss of 0.448, validation loss of 0.4418, and test loss of 0.4379, indicating that it generalizes well to unseen data (see Figure 5). The RMSE is calculated as the square root of the loss function ℒ:

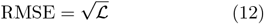

### 5.2 Classification Metrics

The model’s ability to classify secondary structures (*α*-helices and *β*-sheets) was evaluated using precision, recall, F1-score, ROC AUC, PR AUC, and a confusion matrix (see Figure 3). Predictions were binarized using a threshold of 0.3, prioritizing recall for the *β*-sheet class. This threshold was chosen over the default 0.5 due to the structural variability and lower frequency of *β*-sheets. Lowering the threshold improves sensitivity, capturing more true positives for sheets while slightly reducing precision — a trade-off acceptable for accurately identifying sheets in downstream applications.

**Figure 3:**
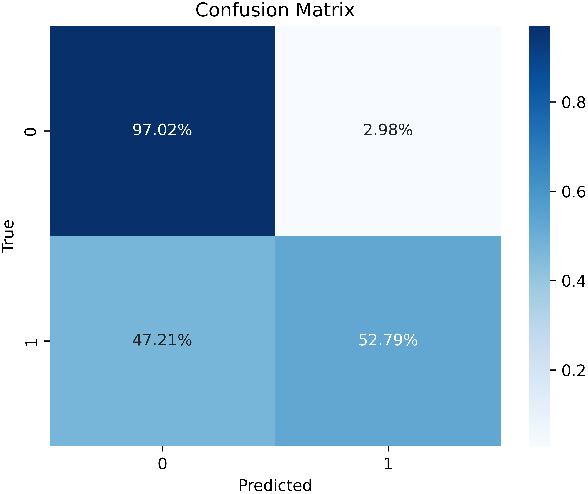
Normalized confusion matrix for secondary structure predictions. Rows represent true labels, and columns represent predicted labels.

The classification metrics are summarized in Table 1. The model achieved high precision (0.9382) and improved recall (0.5279) following threshold adjustment, resulting in an F1-score of 0.6756. The ROC AUC (0.8997) and PR AUC (0.8966) also indicate strong overall performance.

**Table 1:** Classification Metrics for Secondary Structure Prediction.

| Metric | Helix | Sheet | Overall |
| --- | --- | --- | --- |
| Precision | 0.71 | 0.94 | 0.9382 |
| Recall | 0.97 | 0.53 | 0.5279 |
| F1-Score | 0.82 | 0.68 | 0.6756 |
| Accuracy | - | - | 0.77 |
| ROC AUC | - | - | 0.8997 |
| PR AUC | - | - | 0.8966 |

Figure 3 presents the normalized confusion matrix. The model demonstrates a high true positive rate for helices (97.02%), though it struggles to recall sheets (52.79%), which is consistent with the variability of *β*-sheet structures.

These results suggest that the model excels at identifying helices while maintaining high precision for sheets. However, future work should aim to improve sheet recall by leveraging additional structural or evolutionary features.

### 5.3 Qualitative Analysis

In addition to the quantitative evaluation using RMSE values, we qualitatively analyze the model’s performance by visually comparing the predicted protein structures to their corresponding experimentally determined structures using molecular visualization tools like Jalview and PyMOL (see Figure 4). This visualization allows us to observe the local and global conformations of the predicted protein structures and identify regions where the model’s predictions closely resemble or diverge from the true structures.

**Figure 4:**
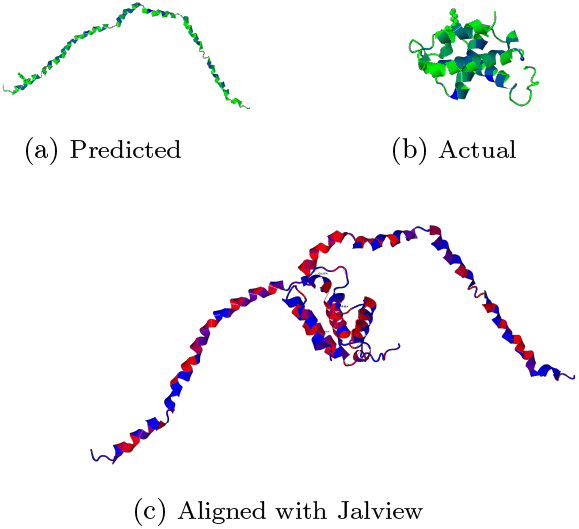
Sample structure prediction on unseen data compared to the actual structure (from ProteinNet) and their alignment

### 5.4 Training Dynamics

The training history over 25 epochs with batch size 4 is shown in Figure 5. The loss curves indicate stable convergence without significant overfitting, except for a noted overfit at epoch 19 where RMSE = ∞.

**Figure 5:**
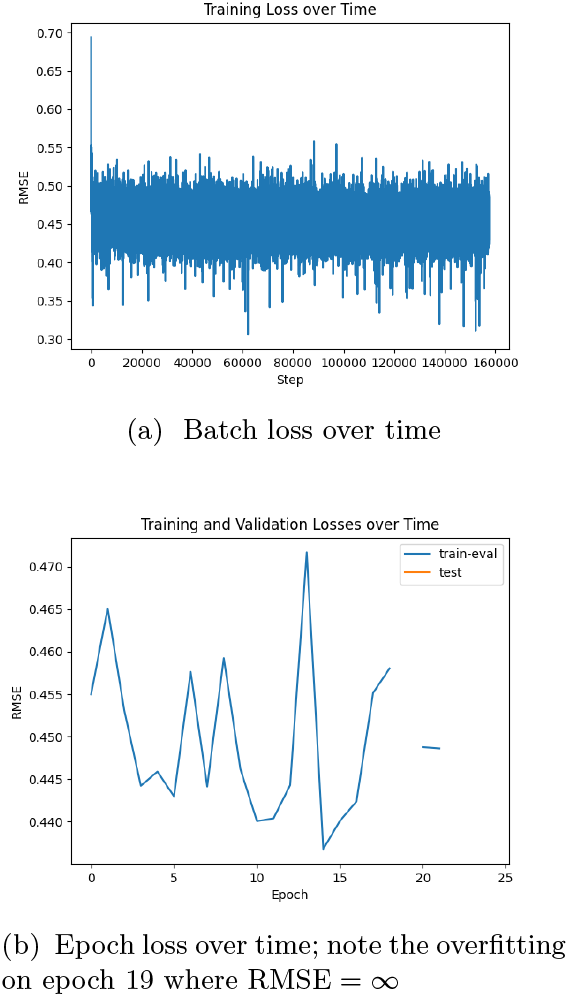
Training history (RMSE loss) over 25 epochs with batch size 4

**Figure 6:**
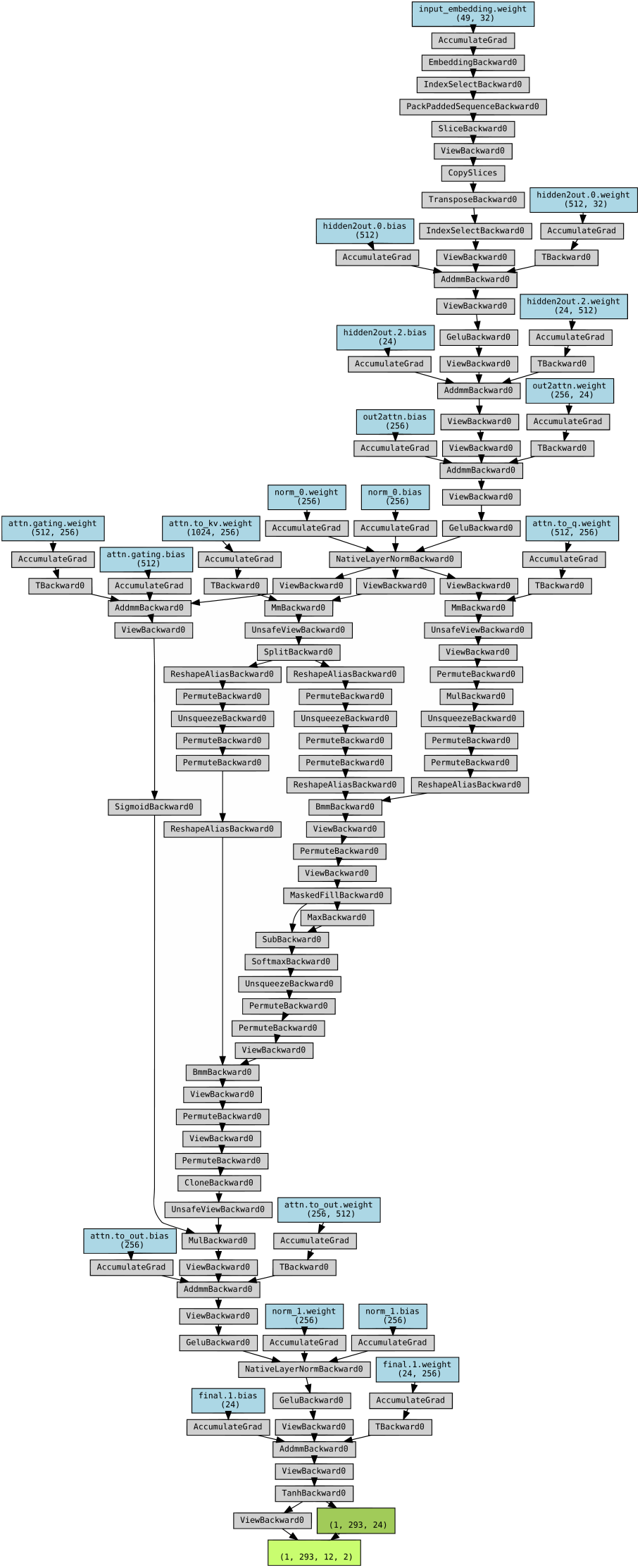
System Architecture Diagram [9]

### 5.5 Secondary Structure Prediction

The model excels in predicting secondary structures, such as *α*-helices and *β*-sheets, which form the local backbone conformation of proteins and are crucial for understanding protein function and stability.

## 6 Discussion

In light of the results obtained from our evaluation, we observe that the model demonstrates promising performance in predicting secondary structures. The RMSE values and classification metrics indicate good generalization to unseen data, with high recall for *α*-helices (97%) and strong precision across all predictions (93.82%). These results indicate good generalization to unseen data and highlight its potential for accurately predicting local backbone conformations of proteins. While we originally attempted to implement the model with TensorFlow and Keras using atomic coordinates as training data, the results indicated drastic overfitting. This initial approach required additional preprocessing and led to increased VRAM requirements during training, yielding suboptimal results.

Despite these successes, the model faces challenges in predicting tertiary structures (which represent the overall folding and 3D arrangement of the protein’s secondary structural elements), primarily due to difficulties in recalling *β*-sheets (52.79%). This limitation reflects the inherent variability and complexity of sheet structures, which often form intricate hydrogen-bonded networks critical for tertiary folds [12]. However, accurate prediction of these structures is essential for deciphering protein-protein interactions, protein-nucleic acid interactions, and the design of novel therapeutics [13].

Studies by researchers such as Röbbe Wünschiers [14], Süreyya Özöğür-Akyüz [15], and Erik Kropat [16] have highlighted the importance of gene networks and their influence on protein expression and folding. Incorporating gene network data into protein structure prediction models could enhance our understanding of the folding process and improve prediction accuracy. By integrating insights from gene regulatory networks, we can potentially capture the complex interactions that influence protein structures.

Furthermore, there is significant interest in understanding the role of protein misfolding and aggregation in neurodegenerative diseases such as Alzheimer’s disease. Research by Ross et al. has investigated the mechanisms of protein aggregation and its implications for neuronal dysfunction [17]. Protein aggregation, or “sticking,” is a critical factor in the pathogenesis of such diseases. By improving our model’s ability to predict tertiary structures and aggregation-prone regions, we can contribute to the early detection and potential therapeutic targeting of neurodegenerative disorders.

Our data analysis methods primarily utilized the Transformer’s attention mechanisms to capture sequence dependencies. To make these methods more explicit and future-oriented, we acknowledge the need to incorporate advanced techniques such as **Graph Neural Networks (GNNs)** and geometric deep learning. These approaches can model the protein structure as a graph, where nodes represent amino acids and edges represent spatial or sequential relationships. By doing so, we can more effectively capture the complex three-dimensional interactions that govern protein folding and function.

To address the identified limitations, future work should focus on expanding the training dataset, possibly by including more diverse protein sequences and structures. Incorporating additional sources of information such as **Multiple Sequence Alignments (MSAs)**, evolutionary coupling data, and gene network data can provide a richer context for the model. Refining the model architecture to include components that capture long-range interactions, such as incorporating recurrent connections or attention mechanisms that consider global context, may improve the model’s ability to predict tertiary structures.

It is important to note that this work is not intended to present a state-of-the-art model, but rather to demonstrate a proof-of-concept using accessible transformer architectures and open-source data. The reliance on CASP12 via SideChainNet was due to its integration and compatibility with our model’s angle-based training pipeline. While newer CASP datasets exist (i.e., CASP15 - 2022), they are not yet fully supported in SideChainNet, and future versions of ProteiNN may incorporate them. We believe that even with its limitations, ProteiNN provides a meaningful contribution as a reproducible and pedagogical benchmark in the growing space of interpretable protein structure prediction tools.

## 7 Conclusion

Our Transformer-based model for protein structure prediction has demonstrated effectiveness in predicting secondary structures, providing a valuable tool for understanding protein function and stability. However, challenges remain in predicting tertiary and quaternary structures, which are critical for fully understanding protein behavior and interactions. Future research should thus focus on several key areas:

1. **Incorporating Gene Network Data**: Integrating gene regulatory network information may enhance the model’s ability to predict protein folding patterns influenced by gene expression dynamics [14].
2. **Addressing Protein Aggregation**: Future models could aim to predict aggregation-prone regions within proteins, contributing to our understanding of diseases like Alzheimer’s where protein aggregation plays a significant role [18].
3. **Utilizing Advanced Modeling Techniques**: Employing graph neural networks and geometric deep learning approaches can better capture the spatial relationships within proteins, potentially improving tertiary structure prediction [19].
4. **Expanding the Dataset**: Including more diverse proteins and leveraging data from MSAs and evolutionary couplings can provide richer training data, enabling the model to generalize better to various protein families [20].
5. **Refining the Model Architecture**: Enhancing the neural network architecture to capture long-range interactions, possibly through attention mechanisms that consider the entire protein sequence or incorporating hierarchical models, can improve prediction accuracy [21].
6. **Exploring Applications in Neuroscience**: Given the connections between protein misfolding and neurological diseases, extending our model to predict structures of proteins involved in such conditions could have significant biomedical implications [22].

By pursuing these directions, we anticipate that our work will contribute to the development of more accurate and comprehensive models for protein structure prediction. This progress is essential for advancing structural bioinformatics, understanding disease mechanisms, and facilitating drug discovery and protein engineering.

## Author Contributions

The author confirms sole responsibility for this work. The author approves of this work and takes responsibility for its integrity.

## Data Availability

The source code of ProteiNN is available at https://github.com/danielathome19/ProteiNN-Structure-Predictor.

## Institutional Review Board Statement

Not applicable.

## Informed Consent Statement

Not applicable.

